# A p32 family RNA editing factor acts in mitochondrial ribosome biogenesis

**DOI:** 10.64898/2026.08.02.742330

**Authors:** Prashant Chauhan, Ingrid Sveráková-Škodová, Jonathan E. Wong, Jan Říha, Julie Dedková, Anna Huber, Lukas Müller, Evgeny S. Gerasimov, Alena Zíková, Vyacheslav Yurchenko, Ondřej Gahura

## Abstract

Biogenesis of mitochondrial ribosomes (mitoribosomes) in the unicellular parasite *Trypanosoma brucei* requires an exceptionally large toolkit of assembly factors, identified in stable precursors of large and small mitoribosomal subunits (mtLSU and mtSSU) by cryoEM. Here, using genetic modifications and proteomic characterization of the immunoprecipitated assemblosome, the earliest characterized mtSSU precursor, we determined that a cap of its distinctive protrusion of hitherto unknown composition consists of a p22 homotrimer. This protein was previously implicated in the uridine-insertion editing of the cytochrome *c* oxidase subunit II transcript. Our functional analysis confirmed this role but revealed that its ablation also causes a loss of mtSSU and a systemic reduction in mitochondrial translation, phenocopying the depletion of established mitoribosomal assembly factors. Consequently, the oxidative phosphorylation system and mitochondrial function are compromised. The p22 protein belongs to the p32 family. We showed that five of its six trypanosomal members are involved in mtSSU biogenesis. Notably, p32 proteins are associated with mitoribosomes in two other distant eukaryotic lineages. A eukaryote-wide mapping of p32 proteins documented that their presence correlates with the retention of mitochondrial genomes. Together, our findings redefine trypanosomal p22 as a dual-function coordinator of mitochondrial gene expression and reveal that the ancestral role of the p32 family is associated with mitochondrial translation.

## INTRODUCTION

Mitochondria are endosymbiotic organelles that have retained reduced genomes and molecular apparatus for their expression. Gene expression processes in mitochondria are frequently elaborated, despite - or perhaps as a consequence of - these genome reductions. Numerous unrelated organisms have independently evolved additional steps in their gene expression pathways, most of which involve complex post-transcriptional RNA processing steps (1).

One of the most striking forms of RNA processing takes place in the mitochondria of kinetoplastid parasites. Approximately two-thirds of the raw protein-coding transcripts in these species undergo site-specific insertion or deletion of uridines to generate mature mRNAs containing functional open reading frames. The positions of these uridine insertions and deletions (U-indels) in all but one of these transcripts are determined by *trans*-acting short guide RNA (gRNA) template molecules encoded within the mitochondrial genome. This entire process, known as kinetoplastid U-indel RNA editing, is mediated by a complex macromolecular machinery comprising several modular and dynamic ribonucleoprotein complexes (2–4).

Mature mRNAs are translated into proteins by mitochondrial ribosomes (mitoribosomes), which have evolved structural features that diverge significantly from those of their bacterial ancestors (5,6). Mitoribosomes are characterized by a high protein-to-RNA ratio, in most eukaryotes contributed by reduction in total RNA content and the acquisition of novel protein elements, lineage-specific mitoribosomal proteins or extensions of proteins of bacterial origin (7,8). This evolutionary trend is particularly pronounced in the mitoribosomes of kinetoplastid parasites (9–11) and their free-living sister lineage, the diplonemids (12). Cryo-electron microscopy (cryo-EM) structures of the mitoribosome from *Trypanosoma brucei* have revealed how this expanded protein repertoire stabilizes highly truncated rRNA with a minimum of double-stranded regions and forms a massive outer shield that cradles the catalytic cores of both the large and small mitoribosomal subunits (mtLSU and mtSSU; (9).

The biogenesis of mitoribosomes is an intricate process, which requires tight coordination between the folding of mitochondrial ribosomal RNAs (rRNAs), encoded in the mitochondrial genome across all eukaryotes, and the sequential binding of ribosomal proteins, a subset of which (or all, in some organisms) is encoded in the nuclear genome and imported from the cytosol. This process is facilitated by assembly factors, which assist in RNA folding, introduce specific RNA modifications, orchestrate protein recruitment, or prevent the premature association of mitoribosomal subunits (8,13–16). Remarkably, the assembly toolkit in *T. brucei* comprises at least 68 assembly factors characterized by cryo-EM studies of precursor mtSSU (17–19) and mtLSU (10,20,21) complexes (pre- mtSSU and pre-mtLSU) (8), considerably outnumbering the assembly factors documented in mammals, which represent the best-understood system of mitoribosome biogenesis.

A large subset of these trypanosomal assembly factors appears to be lineage-specific (18,21), with no known homologs acting in ribosome biogenesis in bacteria or mitochondria of other eukaryotes. Presumably, these factors specialize in stabilizing rRNAs with reduced base-pairing while contributing to the recruitment of the expanded protein complement. Furthermore, unlike in mammals, trypanosomal mitoribosomal precursors are highly abundant (18,20,21), suggesting that some of these assembly factors might mediate functional crosstalk with other mitochondrial processes, possibly aiding in the overall orchestration of mitochondrial gene expression and metabolism (22,23).

Among three trypanosomal pre-mtSSU complexes characterized by cryo-EM, the earliest intermediate, termed the mtSSU assemblosome, is a massive 4 MDa complex containing rRNA in an immature conformation, mitoribosomal proteins, and 35 distinct assembly factors (18,19). A prominent architectural feature of this structure is a tower-like protrusion on the immature intersubunit interface composed of a homopentamer of the assembly factor mt-SAF24, which is capped by a distinct structural density of previously unknown composition. The buried region of mt-SAF24 directly stabilizes an immature conformation of the decoding center (18), suggesting a key role in controlling its maturation.

In this study, we identified the composition of the assemblosome protrusion cap and demonstrated that it is formed by a stable homotrimer of p22, an ortholog of human and budding yeast p32/Mam33, which was previously implicated in *cis*-guided RNA editing of *COII* transcript (24). We show that p22 is essential not only for *COII cis*-editing, but also for the biogenesis of the mtSSU and, consequently, for efficient mitochondrial translation. Mapping of the phylogenetic distribution of p32 homologs revealed that the p32 family is of eukaryotic origin and was present in the last eukaryotic common ancestor (LECA), followed by a multi-paralog expansion in trypanosomatids and plants. The association of p32 with mitoribosomes or their biogenesis across three distant eukaryotic lineages, alongside the absolute correlation between p32 retention and the presence of a mitochondrial genome, strongly implies that the ancestral function of this family is associated with mitochondrial gene expression, most likely translation.

## MATERIALS AND METHODS

### Generation of plasmid constructs and DNA fragments for RNAi, ectopic expression, in situ tagging and conditional knock-out

Regions of TbRbfA (Tb927.11.10600), mt-SAF16 (Tb927.11.3670) and p22 (Tb927.6.2420) for RNAi targeting were selected using RNAit (25). The respective sequences were amplified by PCR using *T. brucei* genomic DNA as a template, KOD DNA polymerase, and primers with extensions for cloning (Table S1). PCR products for RNAi of TbRbfA and mt-SAF16 were inserted into plasmid pTrypSon by Gibson assembly (26) with a *Hind*III-linearized vector backbone and an *Xho*I-digested stuffer fragment using the Gibson Assembly Mix. The PCR product for RNAi of p22 was digested with *BamH*I and *Kpn*I, and with *Xho*I and *Hind*III, and the two products were sequentially cloned into multiple cloning sites I and II, respectively, of pAZ055 (27).

A construct for constitutive expression of mtSAF24ΔNTD^rec^:FLAG, comprising an N-terminal mitochondrial targeting sequence from IscU (Tb927.9.11720; first 57 amino acids) fused to a recoded C-terminal domain of mt-SAF24 with a C-terminal FLAG tag (Table S1), was synthesized and cloned into the *Bam*HI and *Hind*III sites of pHD1344 (28). A construct for tetracycline regulatable expression of mt-SAF24:myc was generated by PCR amplification of full mt-SAF24 CDS with primers including restriction sites and reverse primer containing an extension encoding the myc tag (Table S1). The product was cloned into the *BamH*I and *Hind*III sites of pLEW79 (29). All plasmid constructs were verified by Sanger sequencing.

For C-terminal endogenous V5-epitope tagging, PCR amplicons containing the V5 tag and selection cassette were amplified from pPOTv5 (30) with Expand polymerase and gene-specific primers with 80-nt extensions for homology recombination upstream of the stop codon of the respective gene (Table S1).

To knock out both alleles encoding mt-SAF24, two sgRNAs targeting the 5′ and 3′ ends of the CDS were designed using the LeishGEdit tool (31) and DNA templates for their expression were generated by annealing of a specific forward primer with a universal reverse primer (Table S1) followed by extension with Expand polymerase. Repair templates containing a G418 resistance cassette with 24-nt homology arms were PCR-amplified from pHD2164 (32) using gene-specific primers (Table S1).

### Cultivation and genetic modification of *Trypanosoma brucei*

All cell lines were derived from the procyclic form *T. brucei* Lister 427 strain and are listed in Table S1. The parental strain for tetracycline inducible expression (PF SmOx) was generated by transfection of a construct for expression of T7 polymerase (T7 pol) and tetracycline repressor (tetR) (33). The parental strain for knock-out (PF SmOx Cas9) was generated by transfection of a construct pJ1339 for expression T7 pol, tetR and Cas9 (34).

Cells were cultured at 27°C in SDM-79 medium supplemented with penicillin-streptomycin (100 µg/ml), 4mM hemin, and 10% (v/v) fetal bovine serum, and optionally selection antibiotics, with pH adjusted to 7.3. Cells were maintained in densities between 7×10 and 2×10 cells/ml. Expression of regulatable genes and RNAi constructs was induced with 1 µg/ml tetracycline.

For genetic modifications, cells were transfected by electroporation using the BTX ECM 630 instrument. Prior to transfection, plasmids were *Not*I-linearized, and 10–20 µg of plasmids or DNA fragments were ethanol-precipitated and resuspended in sterile milliQ water. For each transfection, 5×10 cells were harvested from mid-logarithmic cultures by centrifugation at 1,300×g at 12°C, washed once in sterile ice-cold CytoMix (25 mM HEPES, 120 mM KCl, 0.15 mM CaCl₂, 10 mM potassium phosphate buffer, pH 7.6; 2 mM EDTA, 5 mM MgCl₂, and 6 mM glucose) and resuspended in 500 µl CytoMix. DNA was combined with the cell suspension in 2 mm electroporation cuvettes, incubated on ice for 5 min and pulsed at 1,600 V, 25 Ω, and 50 µF. Cells were recovered for 16–18 hours in SDM-79 without selection, then supplemented with an equal volume of SDM-79 containing selection antibiotics at 2× concentration. Selection antibiotics were used at the following final concentrations: G418 - 15 µg/ml, hygromycin - 25 µg/ml, phleomycin - 2.5 µg/ml, puromycin - 1.0 µg/ml and blasticidin - 10 µg/ml. Semi-clonal lines were obtained by limiting serial dilution in 24-well plates and expanded into 25 cm² flasks for screening. Selection of knock-out clones was performed in the presence of 1 µg/ml tetracycline.

### Digitonin subcellular fractionation, western blotting and immunodetection

For whole cell lysate preparation, 1×10 cells were harvested at 1,500×g for 10 min at 4°C, the samples were washed in PBS-G (1× phosphate-buffered saline; 5 mM Na₂HPO₄, 5 mM NaH₂PO₄, 145 mM NaCl, 6 mM glucose, pH 7.4) and resuspended in 100 µl 1× PBS. Samples were combined with 50 µl 3× SDS-PAGE sample buffer (6% SDS, 300 mM DTT, 150 mM Tris/HCl pH 6.8, 30% glycerol, 0.02% bromophenol blue) and heated at 97°C for 10 min. For digitonin subcellular fractionation, 1×10 cells were harvested at 1,500×g for 10 min at 4°C, washed twice in PBS-G and resuspended in 500 µl SoTE buffer (20 mM Tris/HCl pH 7.5, 0.6 M sorbitol, 2 mM EDTA). An equal volume of SoTE containing 0.03% (w/v) digitonin was added, samples were incubated on ice for 5 min, and centrifuged at 4,600×g for 3 min at 4°C. The supernatant containing the cytosolic fraction was transferred to a fresh tube. The pellet with the organellar fraction was resuspended in an equivalent volume of 1× PBS pH 7.4. Aliquots from both fractions were mixed with a 3× SDS-PAGE sample buffer and heated at 97°C for 10 min.

Samples containing material from 2×10 cells were resolved by polyacrylamide gel electrophoresis (PAGE) on 4–20% Tris-Glycine Plus and transferred onto methanol-activated polyvinylidene difluoride (PVDF) membrane using transfer buffer (39 mM glycine, 48 mM Tris, 20% methanol) at 90 V for 90 min at 4°C. Membranes were blocked overnight at 4°C in 5% (w/v) skimmed milk in PBS-T (1× PBS, 0.05% Tween-20) and immunodetection was performed with primary and secondary antibodies (Table S1) in 5% skimmed milk in PBS-T for 90 min at room temperature. The membranes were incubated with the Clarity Western ECL substrate and a chemiluminescent signal was detected on a ChemiDoc XRS+ system. Images were densitometrically analyzed using ImageLab (v6.0.1).

### Isolation of crude mitochondria and native electrophoresis

Crude mitochondria were isolated from 2×10 *T. brucei* cells by hypotonic lysis. Cells were harvested at 1,500×g for 10 min at 4°C, washed in NET buffer (150 mM NaCl, 100 mM EDTA, 10 mM Tris-HCl pH 8.0) and disrupted by ten passages through a 25-gauge needle in DTE buffer (1 mM Tris-HCl pH 8.0, 1 mM EDTA). Sucrose was added to the final concentration of 250 mM. Crude mitochondria were pelleted at 15,000×g for 10 min at 4°C and resuspended in STM buffer (250 mM sucrose, 20 mM Tris pH 8.0, 2 mM MgCl₂). Samples were treated with DNase I in the presence of 3 mM MgCl₂ and 0.3 mM CaCl₂ for 30 min on ice; the reaction was quenched by addition of STE buffer (250 mM sucrose, 20 mM Tris pH 8.0, 10 mM EDTA). Mitochondria were recovered at 15,000×g for 10 min at 4°C and washed twice in STE. The final pellet was aliquoted and either used immediately or snap-frozen and stored at −80°C.

For blue native PAGE (BN-PAGE), crude mitochondria from 5×10 cells were resuspended in ice-cold solubilization buffer (50 mM NaCl, 50 mM Bis-Tris/HCl pH 7.0, 2 mM aminocaproic acid, 1 mM EDTA) supplemented with EDTA-free protease inhibitor) and solubilized with 2% (w/v) n-Dodecyl β-D-maltoside for 1 hour on ice. Insoluble material was removed at 16,000×g for 30 min at 4°C and the protein concentration in the clarified lysates was determined by bicinchoninic acid assay using a bovine serum albumin standard curve. Lysates (10 µg total protein each) were supplemented with native loading dye (0.5 M aminocaproic acid, 5% [w/v] Coomassie G-250) and resolved on NativePAGE 3–12% Bis-Tris gels using the cathode (50 mM Bis-Tris/HCl, 50 mM tricine and 0.002% (w/v) Coomassie G-250, pH 6.8) and anode (50 mM Bis-Tris/HCl and 50 mM tricine, pH 6.8) buffers at 80 V until samples entered the gel, and at 100 V for 3.5 hours at 4°C. The gels were equilibrated in 1× SDS running buffer (25 mM Tris, 192 mM glycine, 0.1% SDS) for 10–15 min, rinsed with milliQ water, and equilibrated in transfer buffer. Proteins were transferred onto the PVDF membrane at 90 V for 100 min at 4°C. Membranes and blocked overnight at 4°C in 5% (w/v) skimmed milk in PBS-T and immunodetection was performed as described above.

### RNA isolation and detection by Northern blotting

Total RNA was isolated by the acid guanidinium thiocyanate–phenol–chloroform method. Cells (5×10) were pelleted at 1,500×g for 10 min at 4°C, washed in 1× PBS and lysed in 500 µl of 4 M guanidinium thiocyanate, 25 mM sodium citrate pH 7.0, 0.5% sarcosyl, 0.1 M β-mercaptoethanol. RNA was extracted by sequential addition of 2 M sodium acetate pH 4.0 and phenol:chloroform:isoamyl alcohol 25:24:1. After vortexing and 15 min incubation on ice, phases were separated by centrifugation at 13,500×g for 15 min at 4°C and RNA was precipitated from the aqueous phase with isopropanol, pelleted, washed with 70% ethanol, air-dried, resuspended in milliQ water, and re-extracted with phenol:chloroform:isoamyl alcohol, and dissolved in 100 µl Diethyl pyrocarbonate-treated water.

For Northern blot analysis, 10 µg of RNA was denatured at 65°C for 10 min in 1.5× loading dye (formamide, 37% formaldehyde, 1× 3-(N-Morpholino)propanesulfonic acid (MOPS)) buffer, bromophenol blue, xylene cyanol, 50 µg/ml ethidium bromide) and resolved on a 1.0% agarose formaldehyde gel in 1× MOPS running buffer (40 mM MOPS, 10 mM sodium acetate, 1 mM EDTA, pH 7.0). Resolved RNA was capillary-transferred overnight in 20× SSC (3M NaCl and 300 mM sodium citrate in Milli-Q water, pH 7.0) onto a positively charged nylon membrane and UV cross-linked to the membrane. Membranes were pre-hybridized for 1 h at 48°C in hybridization buffer (5× SSC, 20 mM sodium phosphate pH 7.2, 7% SDS, 1× Denhardt’s solution (0.02% Ficoll 400, 0.02% polyvinylpyrrolidone K90, 0.02% BSA), 1 mg/ml herring sperm DNA). Oligonucleotide probes were 5′-radiolabelled with γ-³²P-ATP using T4 polynucleotide kinase, heat-denatured, and added to the membrane. Hybridization was performed overnight at 48°C. Membranes were rinsed and washed three times for 20 min in 3× SSC, 5% SDS, three times for 20 min in the same solution supplemented with 10× Denhardt’s solution at 48°C, and once for 20 min in 1× SSC, 1% SDS at 48°C. Membranes were exposed to a storage phosphor screen for 24–48 hours and the signal was detected on an Amersham Typhoon 5 phosphorimager. Images were densitometrically analyzed using ImageLab (v6.0.1). For re-probing, membranes were stripped twice in 0.1× SSC, 0.1% SDS at 80°C for 30 min.

### Quantitative reverse-transcription PCR

Total RNA was isolated from 5×10 cells as described above. RNA concentration was determined using a microvolume spectrophotometer and 5 µg total RNA was treated with TURBO DNase kit according to the manufacturer’s instructions to remove genomic DNA. 1 µg of the DNase-treated RNA was used for cDNA synthesis using Quantiscript Reverse Transcriptase kit at 42°C for 35 min. Subsequently, PCR reactions were prepared in a PCR flow box in a total volume of 15 µl containing 5 µl of 1:10 diluted cDNA, 8 µl SYBR Green master mix and 1 µl of forward and reverse primers (10 µM stock concentration; all primers are listed in Table SX), set up in 96-well plates sealed with optical adhesive film. Reactions were run on a QuantStudio 7 Flex instrument and data analyzed using QuantStudio Real-Time PCR Software (v1.7.2). All RNA samples were processed in technical triplicates from the same cDNA preparation. mRNA levels were normalized against reference gene β-tubulin and expressed as relative fold change over the reference sample using the ΔΔCt method (35). The data were analyzed using QuantStudio Real-Time PCR Software (v1.7.2).

### Mitochondrial membrane potential assessment by flow cytometry

Mitochondrial membrane potential (ΔΨm) was assessed using the fluorescent dye tetramethylrhodamine ethyl ester (TMRE). Mid-logarithmic PF cells (5×10) were harvested at 1,500×g for 10 min at room temperature and resuspended in 1 ml fresh culture medium containing 60 nM TMRE. As a control, a reference sample was in parallel treated with 20 µM uncoupler carbonyl cyanide-4-[trifluoromethoxy]phenylhydrazone (FCCP) to collapse ΔΨm. Samples were incubated for 30 min under standard culture conditions, after which 200 µl was transferred into FACS tubes containing 1 ml 1×PBS. Samples were analyzed on a BD FACSymphony A1 flow cytometer. An FSC-A versus SSC-A gate (P1) was defined to select intact cells and exclude debris; a minimum of 10,000 events per sample were acquired. TMRE fluorescence was detected and median fluorescence intensity (MFI) within the P1 gate was extracted using BD FACSDiva software and analyzed in GraphPad Prism. Data represent three independent biological replicates, each with three technical replicates, expressed as means ± standard deviation. Statistical significance was assessed by paired t-test.

### Immunoprecipitation of V5-tagged mitochondrial proteins and mass spectrometry analysis

Dynabeads Protein G were washed twice in 1×PBS, coupled to anti-V5 monoclonal antibodies (2 µg per 15 µl beads) in 1×PBS overnight at 4°C with rotation. Beads were cross-linked with 3 mg/ml dimethyl suberimidate dihydrochloride in 1×PBS pH 8.0 for 45 min at room temperature with rotation; the reaction was quenched with Tris/HCl pH 7.5 (50 mM final) for 15 min at room temperature. Cross-linked beads were washed twice in 1×PBS and twice in IP buffer (25 mM Tris/HCl pH 7.5, 100 mM KCl, 15 mM MgCl₂, 0.1% Triton X-100, EDTA-free protease inhibitor). Four biological replicates were processed in parallel, each from mitochondria from 5×10 cells. Mitochondria were lysed in 100 µl IP buffer with 1.7% (v/v) Triton X-100 for 1 hour on ice. Insoluble material was removed at 15,000×g for 1 hour at 4°C. A 10 µl aliquot of the clarified lysate was saved as input; the remainder was combined with cross-linked beads in a total volume of 200 µl and incubated for 6 hours at 4°C with rotation. The unbound flow-through was retained for western blot analysis. Beads were washed twice in IP buffer and three times in 1×PBS (5 min each, 4°C). The samples were snap-frozen in liquid nitrogen and stored at −80°C for mass spectrometry (MS).

Samples were analyzed by label-free quantification MS following the SP3 protocol with glass beads (36). Proteins were solubilized in 1% SDS/100 mM triethylammonium bicarbonate, reduced with 10 mM Tris(2-carboxyethyl)phosphine (TCEP) and alkylated with 40 mM chloroacetamide (95°C, 10 min). MS-grade trypsin digestion was performed overnight at 37°C (1:40 enzyme-to-protein ratio); peptides were desalted by C18 StageTips and 500 ng per sample separated on a C18 column using a Dionex Ultimate 3000 nanoUHPLC. Samples were acquired in DDA mode on an Orbitrap Exploris 480 equipped with a FAIMS unit and processed in MaxQuant; DIA MS Thermo raw files were additionally processed in Spectronaut (Biognosys) against the *T. brucei* TREU927 proteome (TriTrypDB-68_TbruceiTREU927_AnnotatedProteins.fasta) with default settings and precursor, protein Q-value and PEP cutoffs set at 0.01. Protein group quantities (PG.Quantity, MS2 level) from Spectronaut’s protein report were imported into Perseus (37), log₂-transformed, and filtered to remove reverse hits, contaminants and site-only identifications. Protein groups with fewer than two valid values were excluded; remaining missing values were imputed from a normal distribution (width 0.3, downshift 1.8, per column). Differential enrichment between V5-tagged and untagged conditions was assessed by a two-sided two-sample t-test with permutation-based FDR correction (250 randomizations, FDR 0.05, S0 = 0.1), with the V5-tagged condition designated as the right group, and results visualized as volcano plots. Enrichment thresholds were set at difference log_2_ fold change ±1.58 (3-fold change) and −log_10_(*P* value) > 1.3 (*P* < 0.05).

### Metabolic labeling and electrophoretic analysis of products of mitochondrial translation

Mitochondrial translation products were labelled in vivo as follows. 1×10 cells from a logarithmic culture were harvested at 1,000×g for 5 min at room temperature, washed twice in SoTE, pelleted at 1,100×g for 5 min and resuspended in 90 µl SoTE. Cycloheximide (200 µg/ml) and DTT (2 mM) were added to inhibit cytosolic translation and prevent cell aggregation, and cells were incubated at 27°C for 10 min with shaking. Labelling was initiated by addition of 10 µl [³ S]-Met label mix (Hartmann Analytic, IS-103) per sample and continued for 1 h. Labelled products were resolved by two-dimensional SDS-PAGE (9% first dimension, 14% second dimension) and visualized by autoradiography as described (38). The signal was quantified using ImageJ (1.54g). Total protein was visualized by Coomassie R-250 staining.

### Structure prediction and analyses

Structures of proteins were predicted by AlphaFold 3 (39). Structures were visualized and aligned in ChimeraX (40). The structure of the p22 trimer was fitted to the mtSSU assemblosome density using ChimeraX and adjusted manually. Targeting prediction was carried out with DeepLoc 2.1 (41).

### Analyses of the phylogenetic distribution of p22 homologs

The sequences of *T. brucei* mSAF-16, mSAF-19, mSAF-25 and p22, human p32, and *Saccharomyces cerevisiae* Mam33 were used for BLASTp (42) search in the UniProt database. The retrieved sequences were analyzed by CLANS clustering (43), Swiss-model (44), and HHpred (45) to filter out false hits and aligned by MAFFT v7.525 (L-INS-i) (46) to build a profile hmm using HMMER3 (47). Search using HMMER3 was performed in EukProt v3 TCS (48) (The Comparative Set – 196 selected species across eukaryotic diversity). Sequences were cleaned from contaminants, duplicates and false positive hits with phylogenetic approach - sequences were aligned by MAFFT, trimmed by BMGE v1.12 (49) and phylogenetic trees were constructed in IQ-TREE v3.0.1 (50). Next, we searched with HMMER3 and BLASTp across 29 selected species from Euglenozoa available on EukProt v3, and sequences were filtered as described above. In addition, we used HMMER3 to search 50 bacterial and archaeal genomes obtained from NCBI RefSeq, but identified no p32 homologs.

For the reconstruction of the phylogenetic tree, a structure-based alignment was created by PROMALS3D (51) using experimental structures of human p32, *S. cerevisiae* Mam33, and *T. brucei* mt-SAF16, mt-SAF19, mt-SAF25, as well as *T. brucei* and *Leishmania* sp. p22 (PDB IDs 1P32 (52), 3QV0 (53), 6SGB - chains FH, FK and FH (18), 3JV1 (24) and 1YQF), and AlphaFold3 (39) predicted structures of two remaining *T. brucei* p32 paralogs and homologs from *Paradiplonema papillatum* and *Euglena gracilis*. Other sequences obtained by HMMER3 (47) search were aligned to the seed structural alignment using MAFFT *--add* option with L-INS-i algorithm (46) and manually trimmed. The phylogenetic tree was constructed using IQ-TREE v3.0.1 (50) with the LG+C60+G4 model with statistical support assessed by 1000 ultrafast bootstraps with bnni correction (54) and SH-aLRT test (55).

### Reagents

Lists of all antibodies, oligonucleotides, and reagents and chemicals, including manufacturers and catalogue numbers are provided in Table S1.

### Biological resources

Lists of all plasmid constructs and strains generated in this study are provided in Table S1.

### Statistical analyses

Statistical analyses were performed using GraphPad Prism (v11.0.2); two-tailed paired t-tests were used for two-group comparisons, with significance set at *p* value < 0.05. Samples for quantitative MS were analyzed in three independent biological replicates. MS data were processed using Spectronaut, MaxQuant (v2.8.1.0), and Perseus (v2.1.x), output matrices from Perseus were analyzed in MS Excel and GraphPad Prism (v11.0.2). Genomic and protein sequence information was retrieved from TriTrypDB Release 69 (March 2026 (56)).

## RESULTS

### A precursor of the small mitoribosomal subunit contains the RNA editing factor p22

A structural hallmark of the earliest characterized mtSSU assembly intermediate in *T. brucei*, termed the assemblosome, is a tower-like protrusion on the immature intersubunit side (18). The protrusion is constituted by the N-terminal domains (NTD) of five copies of the assembly factor mt-SAF24 and is capped with a distinct disc-shaped structure of unknown composition. The C-terminal domain (CTD) of the homopentameric mt-SAF24 is buried in the assemblosome and keeps mtSSU ribosomal RNA (rRNA) helices h18 and h44, which are essential for mRNA decoding, in an immature conformation (Fig. 1A).

**Figure 1.**
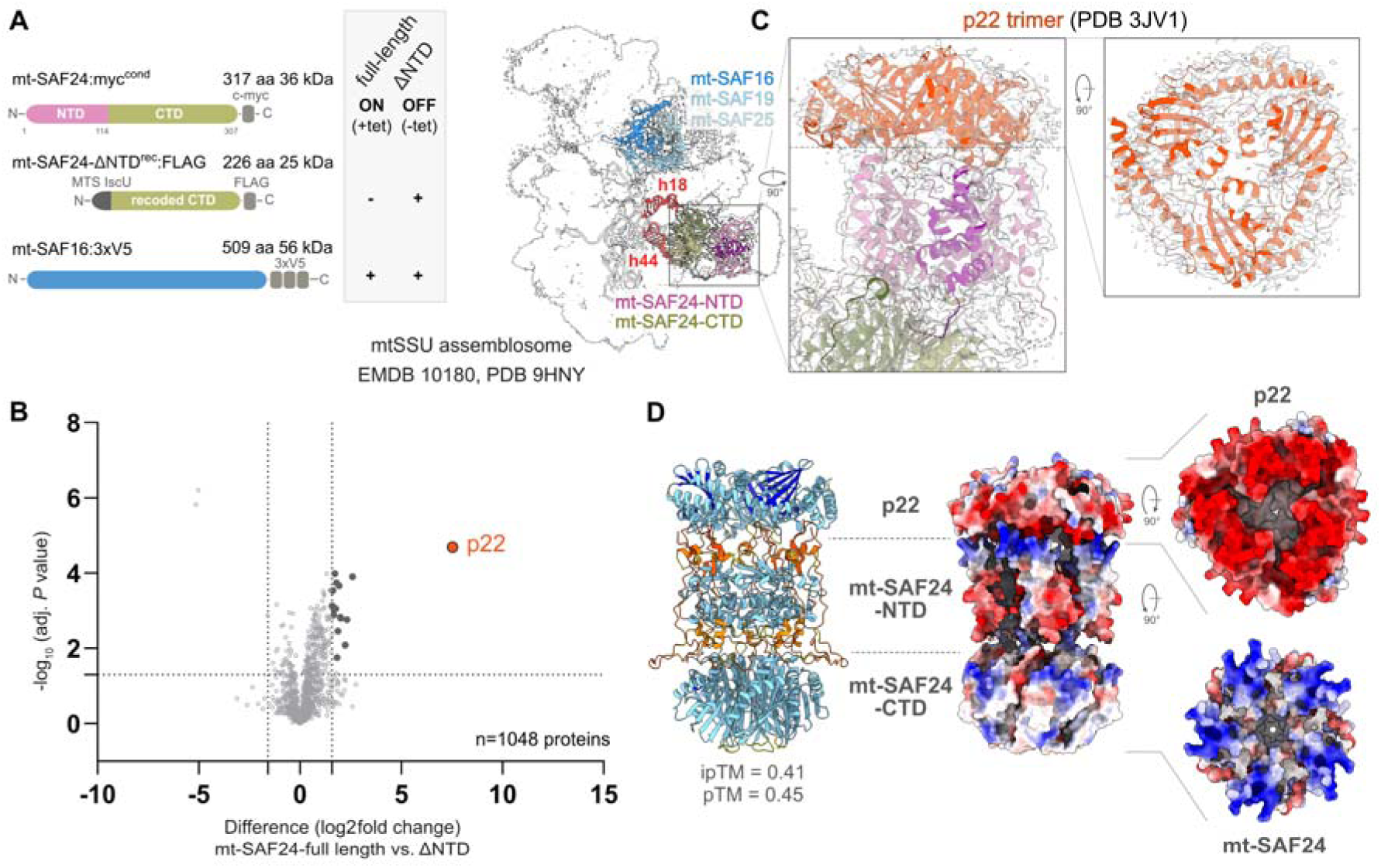
An early precursor of mtSSU in *Trypanosoma brucei* contains a trimer of p22. **(A)** Scheme of genetic modifications in the cell line used for the determination of the unassigned density content (left) and the structure of mtSSU assemblosome show as transparent cryoEM density map (EMDB 10180 (18)) with proteins of interest and rRNA depicted as ribbon (PDB 9HNY (19); right). **(B)** Dot plot showing a comparison of proteins immunoprecipitated by mt-SAF16:3xV5 protein from cells expressing full-length mt-SAF24 and its truncated mt-SAF24-ΔNTD^rec^:FLAG variant, revealing the compositional difference between the cap-containing and cap-deficient complex. **(C)** The crystal structure of p22 (PDB 3JV1 (24)) fitted into the mtSSU assemblosome density. **(D)** AlphaFold model of the assembly of pentameric mt-SAF24 and trimeric p22. The surface colored by Coulombic electrostatic potential (charge) is shown for side view and for the interacting interfaces.

To identify the protein composition of the cap, we genetically modified the procyclic form of *T. brucei* to remove the NTD of mt-SAF24, thereby decoupling the cap from the remainder of the complex. First, we generated a conditional mt-SAF24 knock-out strain (mt-SAF24^cKO^) by introducing a tetracycline-inducible myc-tagged ectopic mt-SAF24 allele (mt-SAF24:myc^cond^) followed by CRISPR-Cas9 driven deletion of both endogenous mt-SAF24 alleles (Fig. S1A-D). The cell line reproduced a strong growth defect and decrease in mtSSU rRNA levels associated with the ablation of mt-SAF24 (Fig. S1E,F) that was reported previously (18). Next, we introduced a constitutively expressed FLAG-tagged truncated version of mt-SAF24 without NTD (mt-SAF24-ΔNTD^rec^:FLAG) and verified its targeting to mitochondria by digitonin-based subcellular fractionation (Fig. S1G,H). Finally, we endogenously tagged one allele encoding assembly factor mt-SAF16 with a 3×V5 epitope (mt-SAF16:3×V5) to enable immunoprecipitation (IP) of the assemblosome (Fig. 1A, S1H).

Using label-free mass spectrometry, we compared the protein content between complexes with and without the cap immunoprecipitated from mitochondria isolated from cells expressing full-length mt-SAF24:myc^cond^ and truncated mt-SAF24-ΔNTD^rec^:FLAG, respectively. The comparison pointed to a single candidate for the unassigned cap density, p22 (TriTryp ID Tb927.6.2420, Uniprot ID Q584R4; Fig. 1B; Table S1), a protein previously implicated in U-indel RNA editing in *T. brucei* mitochondria (24,57,58).

The crystal structure of a p22 homotrimer was resolved by X-ray crystallography (24) (PDB 3JV1). Its atomic model fits into the cryoEM density of the pre-mtSSU protrusion cap (Fig. 1C). Furthermore, structural predictions by AlphaFold support a direct physical interaction between mt-SAF24 and p22 in a configuration that is fully compatible with the experimental assemblosome cryo-EM map. The contacts between the two proteins are provided predominantly by extensive polar interactions between the highly negatively charged surface of the p22 trimer and a complementary positively charged surface on the mt-SAF24 pentamer (Fig. 1D). Together, these biochemical and structural data document that the capping density of the pre-mtSSU protrusion is formed by a homotrimer of p22.

### p22 interacts with RNA editing substrate binding complex subunits and is required for editing of *COII*

In *Trypanosoma cruzi*, p22 was proposed to regulate the abundance of cytoplasmic mRNA-containing granules (59). However, the cytosolic fraction of *T. brucei* lysates obtained by digitonin fractionation contained no detectable levels of p22. The protein is found exclusively in the organellar fraction (Fig. 2A), implying mitochondrial localization and function. Prior studies documented that in *T. brucei* p22 interacts with RBP16 (57), and KREL1 and RESC13/RGG2 (24), proteins acting in kinetoplastid U-indel RNA editing.

**Figure 2.**
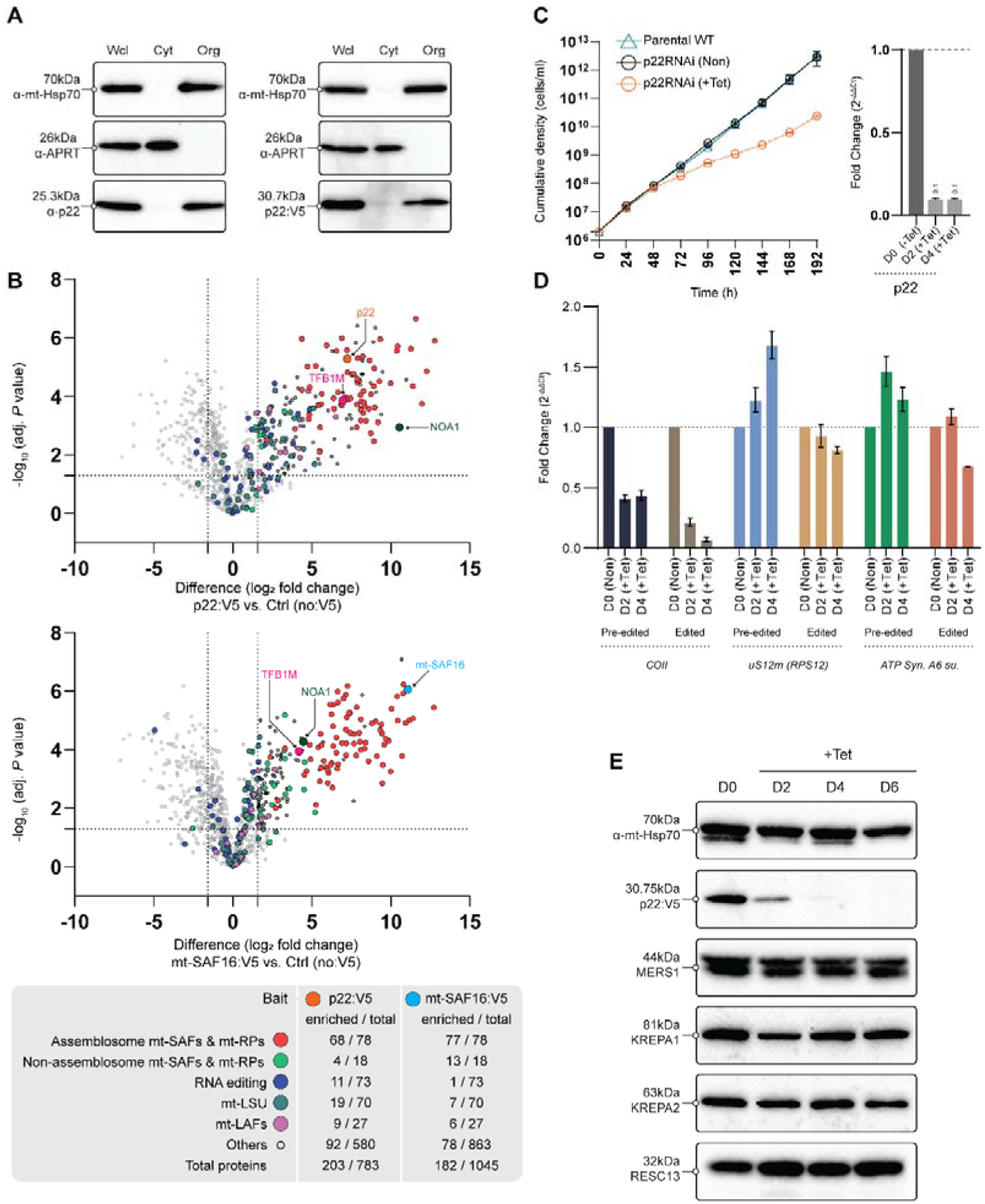
p22 interacts with components of the RNA editing machinery and is required for efficient *COII* editing. **(A)** Immunoblotting of whole-cell lysates (Wcl), and cytosolic (Cyt) and organellar (Org) fractions with anti-p22 and anti-V5 antibodies. **(B)** Dot plots showing proteins detected by MS after IP with anti-V5 antibody from cells expressing p22:V5 and mt-SAF16:V5 and summary of enriched and total number proteins from indicated classes. **(C)** Growth curve of p22 RNAi cells and p22 mRNA levels quantified by qRT-PCR prior and after induction of p22 RNAi. **(D)** Levels of indicated pre-edited and edited mRNA in the p22 RNAi cell line determined by qRT-PCR. **(E)** Impact of p22 RNAi on the steady-state levels of proteins acting in RNA editing and processing assayed by immunoblotting.

To map the interactome of p22, we endogenously tagged the proteins with a V5-epitope and identified its interaction partners by IP followed by label free-MS (Table S1). This analysis confirmed that p22 stably associates with the mtSSU biogenesis machinery, as 67 out of 79 assemblosome components were significantly enriched (at least 3-fold enrichment; Fig. 2B). A total of 11 proteins belonging to RNA editing and processing machinery (3) were also detected as p22 interactors, although their enrichment was lower than that of assemblosome subunits (Fig. 2B). Among these, seven proteins co-constitute the RNA editing substrate binding complex (RESC). The remaining identified proteins were KPAF1, KPAF4 and MERS1, which act in 5’- and 3’-end RNA processing, and the RNA editing helicase KREH2 (3). We also identified a minor subset of mature mtLSU proteins (19) and assembly factors (9).

For comparison, we used the same IP-MS approach to define the interaction network of an established mtSSU assembly factor, mt-SAF16 (Table S1). Like p22, mt-SAF16 co-immunoprecipitated nearly the entire assemblosome (77 enriched components, including p22) and, in addition, previously unidentified homologs of the human mtSSU assembly factors NOA1 (Tb927.6.2290) and TFB1M (Tb927.10.3690; (60–62)Fig. S2). However, with mt-SAF16:V5 the only enriched protein acting in RNA editing was RNA-editing 3 terminal uridylyl transferase 1 (KRET1). This observation indicates that the link to RNA editing is not a common feature of the pre-mtSSU complex, rather it is a specific feature of p22.

Previously, the downregulation of p22 was shown to selectively impair the editing of *COII* (24), which encodes a core subunit of cytochrome *c* oxidase (or complex IV; cIV), the terminal complex of the electron transport chain. We generated an independent cell line allowing inducible suppression of p22 expression by RNAi. We confirmed that the ablation of p22 to approximately 10 % at the mRNA level, which resulted in nearly undetectable protein levels, caused a severe growth defect (Fig. 2C). Next, we assayed the efficiency of RNA editing for *COII*, *RPS12* (encoding mitoribosomal protein uS12m), and *A6* (encoding subunit a of ATP synthase). In agreement with earlier work (24), the loss of p22 had only a marginal effect on the editing of *RPS12* and *A6*. In contrast, the levels of edited *COII* transcripts decreased to 20% and 4% at two and four days post-RNAi induction, respectively, confirming a profound and specific defect in *COII* editing. Notably, we also observed a previously unreported 60% reduction of pre-edited *COII* transcripts relative to non-induced control cells (Fig. 2D). This drop can be explained either by a decreased stability of the pre-edited RNA substrate in the absence of a protective editing complex, or by a gene-specific defect in an upstream process, such as mitochondrial transcription.

To determine if these changes reflected broader defects in the editing machinery, we assayed the steady-state levels of core subunits from distinct RNA processing complexes, specifically KREPA1 and KREPA2 (components of the RNA editing catalytic complex, RECC), RESC13 (RESC), and MERS1 (3’-end processing module), by immunoblotting of mitochondrial lysates resolved by denaturing electrophoresis. Downregulation of p22 caused no detectable decrease in the abundance of these proteins (Fig. 2E). Together, these data demonstrate that the loss of p22 does not compromise the overall integrity of the mitochondrial RNA editing machinery or alter the global U-indel editing profile. Instead, p22 is selectively required for the processing of a single pre-mRNA transcript (24).

### Oxidative phosphorylation complexes are globally reduced upon p22 knockdown

To assess the impact of p22 depletion on the expression of mitochondrial-encoded subunits, which generally cannot be detected by mass spectrometry in *T. brucei* (63) due to their extreme hydrophobicity, we assayed the levels of oxidative phosphorylation (OXPHOS) complexes of dual genetic origin and their individual subunits using denaturing and native gel electrophoresis followed by immunoblotting. Under denaturing conditions, we observed a decrease in the abundance of the cI subunit NDUFA6 and the cIV subunit COXEG7, whereas the levels of ATP synthase subunits β and OSCP remained largely unaffected (Fig. 3A).

**Figure 3.**
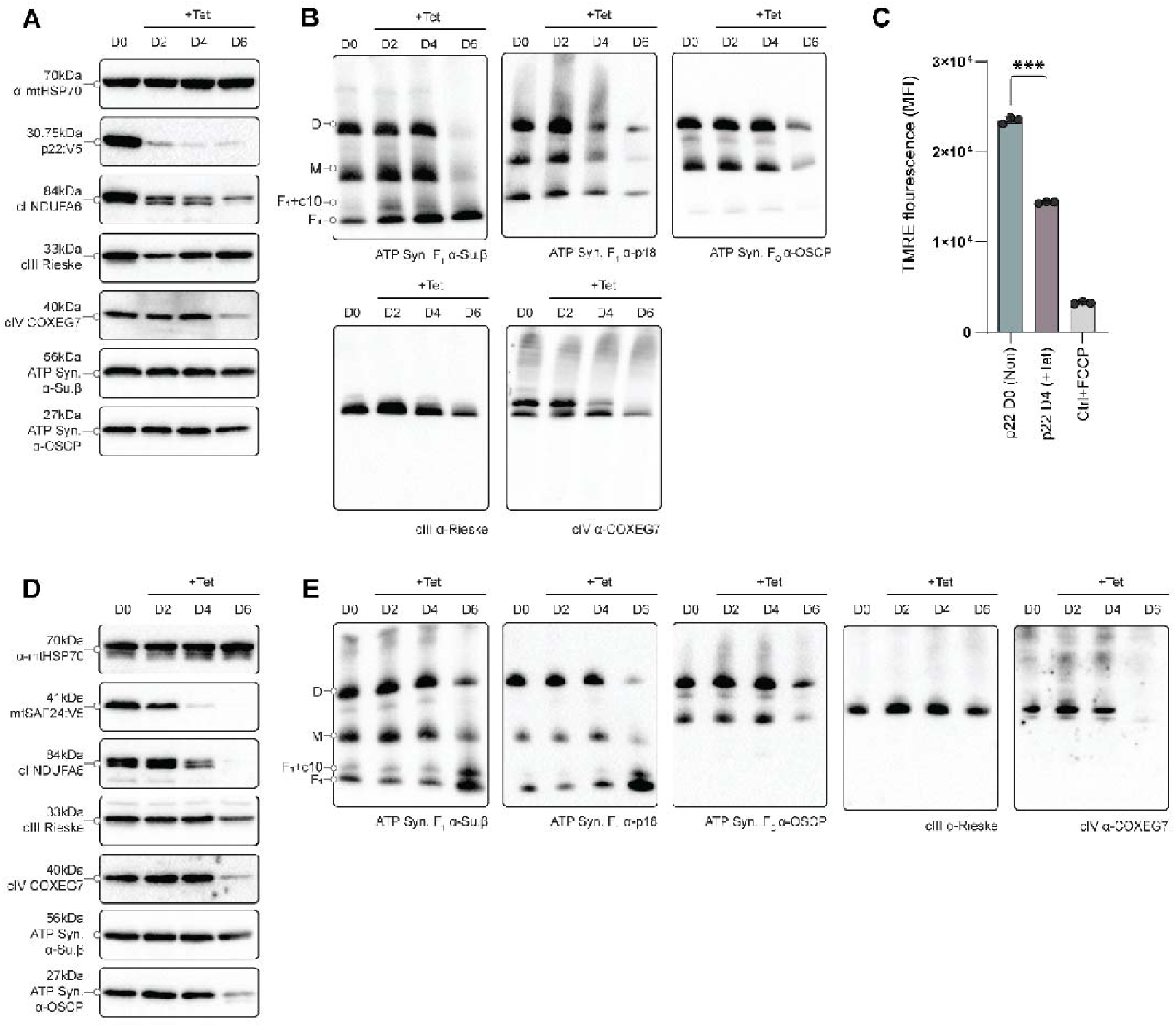
Knockdown of p22 and mt-SAF24 affects the levels of all oxidative phosphorylation complexes containing mitochondrial-encoded subunits and generation of mitochondrial membrane potential. **(A,D)** Steady-state levels of the target V5-tagged proteins and indicated subunits of OXPHOS complexes assayed by immunoblotting of mitochondrial lysates from p22 (A) and mt-SAF24 (D) RNAi cell lines resolved by SDS-PAGE. **(B,E)** Levels of ATP synthase, cIII, and cIV assayed by immunoblotting of mitochondrial lysates from p22 (B) and mt-SAF24 (E) RNAi cell lines resolved native PAGE. **(C)** Changes in mitochondrial membrane potential (Δψ_m_) upon p22 RNAi knockdown measured by TMRE fluorescence. The data points represent averages of three technical replicates from three independent experiments; the error bars represent standard deviation. Statistical significance was assessed by paired t-test (*** *p*<0.0003).

To detect the levels of assembled OXPHOS complexes, mitochondrial lysates were resolved on native gels. Complex III (cIII; ubiquinol-cytochrome *c* oxidoreductase), cIV and ATP synthase were detected by specific antibodies at several time points after p22 RNAi induction. We observed a strong reduction in cIV, consistent with the specific loss of edited *COII* transcripts. However, the abundance of assembled cIII and both the monomeric and dimeric forms of ATP synthase also decreased markedly, albeit at later time points post-induction. The soluble catalytic domain of ATP synthase, F_1_- ATPase, remained largely unaffected, consistent with its independent assembly (27) and the fact that it is composed entirely of nuclear-encoded subunits (Fig. 3B). The membrane-embedded part of ATP synthase contains the mitochondrial encoded subunit-a, whose transcript editing is only slightly affected in p22 RNAi cells ((24) Fig. 2D). The editing of cytochrome B (cyB) transcript, the only mitochondrial encoded subunit of cIII, is not compromised in p22 RNAi cells at all (24). To compare the phenotypes of p22 ablation with those associated with the suppression of an established mtSSU assembly factor, we knocked down mt-SAF24 by RNAi. The pattern of disruption of OXPHOS complexes and their subunits in this cell line resembled that of p22 RNAi. (Fig. 3D,E).

To assess the broader physiological impact of the ablation of p22 on mitochondrial function, we measured the mitochondrial membrane potential (ΔΨ_m_) in living cells using the fluorescent dye TMRE, which accumulates in mitochondria in ΔΨ_m_-dependent manner. Flow cytometry analysis revealed a substantial decrease in ΔΨ_m_, in p22 knockdown cells, indicating a profound electron transport chain deficiency (Fig. 3C,F).

Together, these results demonstrate that the loss of p22 causes defects across multiple OXPHOS complexes containing mitochondrial-encoded subunits, which cannot be explained solely by an inefficient *COII* transcript maturation. Instead, these data indicate that the phenotypes stem from a role for p22 during mtSSU biogenesis, a conclusion supported by the phenocopying observed upon knockdown of the assembly factor mt-SAF24.

### Ablation of p22 impairs mtSSU abundance and causes mitochondrial translation defects

The downregulation of established mtSSU biogenesis factors in trypanosomes typically results in severe growth retardation accompanied by a selective decrease in the levels of mtSSU rRNA, leaving the mtLSU rRNA largely unaffected (18). We confirmed this characteristic phenotype upon RNAi-mediated knockdown of two other core mtSSU assembly factors: TbRbfA (mt-SAF18) and mt-SAF16 (Fig. 4A,B). The downregulation of mt-SAF16 also resulted in decreased steady-state levels of mitoribosomal protein mS64 endogenously tagged with a V5-epitope (Fig. 4C). Because m64 is recruited to the mtSSU in a post-assemblosome stage of maturation (8), the reduction of its levels indicates that upstream assembly steps have been compromised.

**Figure 4.**
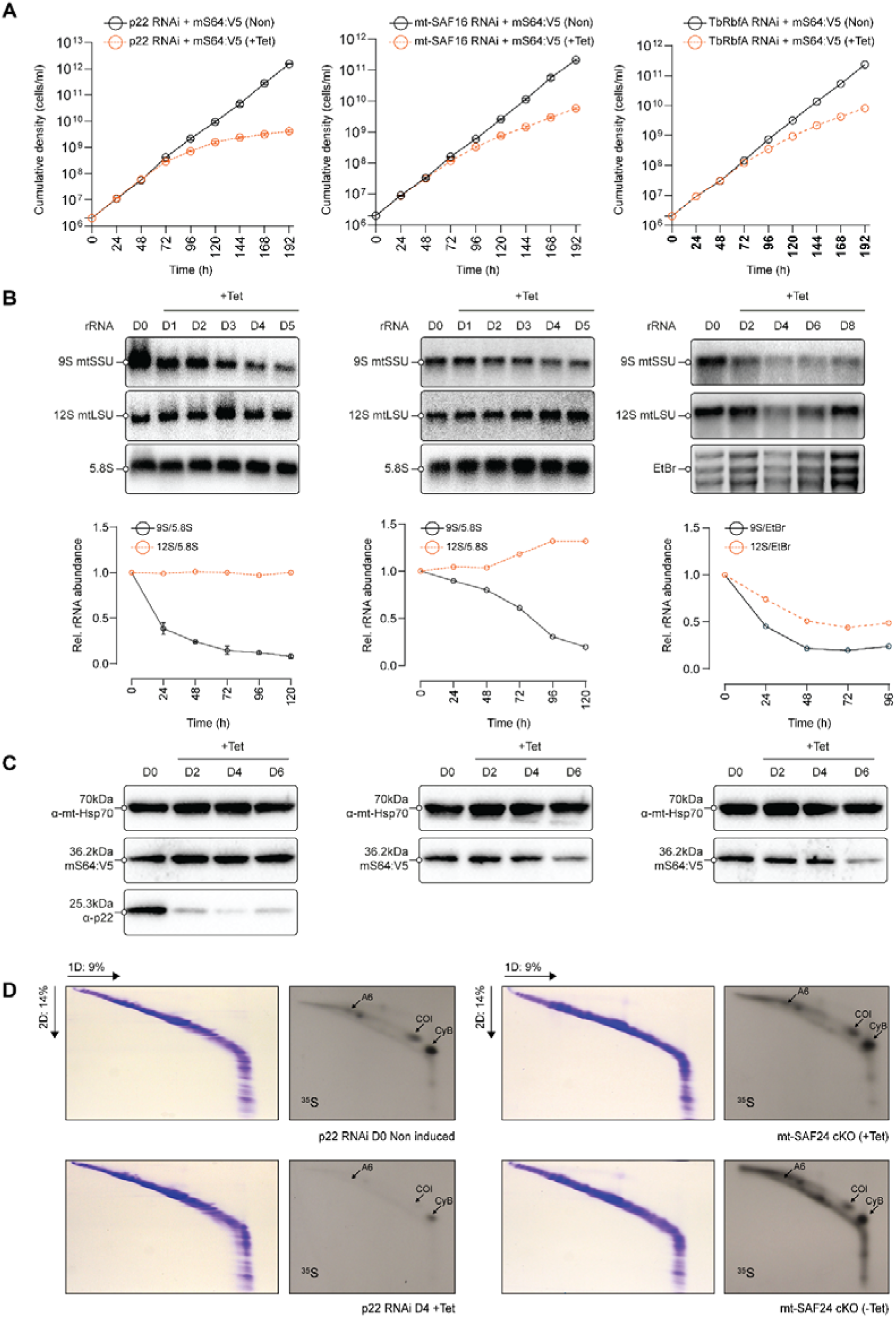
Levels of mtSSU and mitochondrial translation efficiency are reduced upon knockdown of p22 and other assembly factors. **(A)** Growth curves of non-induced and tetracycline-induced p22, mt-SAF16 and TbRbfA RNAi cell lines with V5-tagged mS64. **(B)** Northern blot analysis of mtSSU (9S) and mtLSU (12S) rRNA levels after induction of p22, mt-SAF16 and TbRbfA RNAi. The levels of cytosolic 5.8S rRNA were or a signal from ethidium bromide-stained gel prior transfer were used as a reference for quantification shown in the lower panel. **(C)** Levels of mS64:V5 after p22, mt-SAF16 and TbRbfA RNAi induction assayed by immunoblotting. Probing of mt-Hsp70 was used as a loading control. **(D)** Products of ^35^S metabolic labelling of products of mitochondrial protein translation from non-induced and tetracycline-induced p22 RNAi cells and mt-SAF24^cKO^ strain with and without tetracycline resolved by two-dimensional SDS-PAGE and detected by autoradiography. Coomassie stained gels are shown to document equal loading. Proteins produced by mitoribosome are marked.

We next tested whether p22 ablation leads to a similar mtSSU assembly defect. Indeed, the mtSSU rRNA was markedly reduced at early time points following p22 RNAi induction (Fig. 4A,B). However, the steady-state protein levels of mS64 were not altered by p22 RNAi (Fig. 4C). This distinct phenotype implies that while p22 ablation compromises mtSSU assembly, the structural lesion or assembly block differs from that induced by the loss of mt-SAF16 or TbRbfA.

To test whether the decreased levels of mtSSU rRNA directly impacts organellar protein synthesis, we pulse-labeled newly synthesized, mitochondrially encoded proteins using ^35^S-methionine/cysteine. The labeled translation products were resolved on two-dimensional SDS-PAGE gels using an established procedure (64) to separate and detect labeled cytochrome b, COI, and subunit a, components of cIII, cIV and ATP synthase, respectively. The de novo synthesis signals corresponding to all three detectable mitochondrial encoded proteins were reduced upon the downregulation of p22 (Fig. 4C), demonstrating that p22 is required for efficient mitochondrial protein synthesis. The intensity of CyB and COI signals was reduced by 52 and 61% compared to uninduced cells. An analogous, yet weaker (6% and 33% decrease), translation defect was observed when the essential assembly factor mt-SAF24 was knocked out (Fig. 4C).

Collectively, these results show that p22 is required to maintain physiological pools of functional mtSSUs and is therefore indispensable for efficient mitochondrial translation.

### An expanded p32 family act in mtSSU assembly in *Trypanosoma brucei*

Trypanosomal p22 is a homolog of human p32 (or C1QBP) and *Saccharomyces cerevisiae* Mam33p, and, as such, is a member of the p32 family (57). Mammalian p32 was reported to localize to multiple cellular compartments and participate in diverse processes, including biogenesis of mtSSU (see Discussion). Notably, besides p22 identified in this study, the assemblosome contains additional three proteins belonging to the p32 family - mt-SAF16 (Tb927.11.3670), mt-SAF19 (Tb927.7.7080) and mt-SAF25 (Tb927.10.1820; (19)). In contrast to the homotrimeric configuration of p22, these three paralogs assemble into a distinct mt-SAF16/19/25 heterotrimer, which localizes to a wedge of assembly factors between the head and body of the pre-mtSSU (18,19); Fig. 1A).

We searched the *T. brucei* predicted proteome and found two additional homologs of p32 (TriTrypDB IDs Tb927.7.3470 and Tb927.11.9600, annotated as a p22-like protein and a hypothetical protein, respectively). Although the overall sequence similarity among the six trypanosomal p32 family members is rather low (Fig. S3), a comparison of the crystal structure of the p22 homotrimer and the cryo-EM structure of the mt-SAF16/19/25 heterotrimer with AlphaFold-predicted models of the remaining two paralogs revealed overall conservation of core tertiary structural features (Fig. 5A).

**Figure 5:**
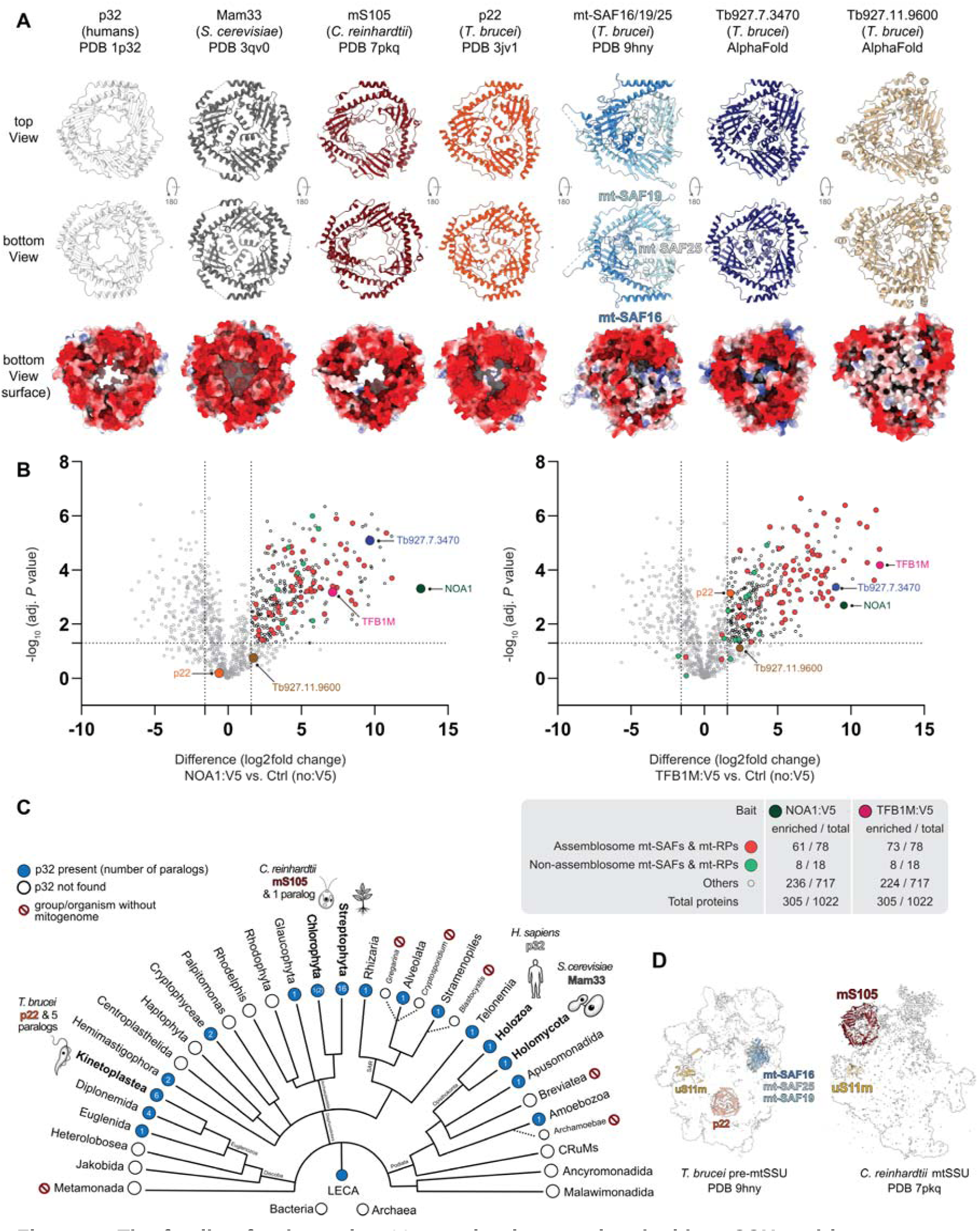
The family of eukaryotic p32 proteins is associated with mtSSU and its precursors and is expanded in trypanosomes. **(A)** Gallery of p32 family proteins and their negatively charged surfaces. **(B)** Dot plots showing proteins detected by MS after IP with anti-V5 antibody from cells expressing NOA1:V5 and TFB1M:V5 and summary of enriched and total number proteins from indicated classes. **(C)** Distribution of p32 family members in eukaryotes including number of paralogs in individual groups. **(D)** Localization of the mt-SAF16/19/25 heterotrimer and p22 homotrimer in *T. brucei* pre-mtSSU and mS105 in *C. reinhardtii* mtSSU. The cryoEM maps are shown as transparent surfaces and the proteins of interest, including uS11m, as ribbons. Both complexes are viewed from the solvent exposed side.

Each protomer across the various trimers consist of a central β-sheet composed of six to seven antiparallel β-strands twisted around two tandem C-terminal α-helices, preceded by an N-terminal helix (which is absent only in mt-SAF25). The same arrangement is observed in trimers of the crystal structures of human p32 and yeast Mam33 (Fig. 5A; (52) (53). The p32/Mam33 proteins are highly acidic, with negatively charged residues distributed asymmetrically on one side of the trimer. The homotrimers of p22 and the uncharacterized paralog Tb927.7.3470 also exhibit extremely negatively charged surfaces on the side, which in case of p22 faces mt-SAF24 (Fig. 1D). Within the mt-SAF16/19/25 heterotrimer, the highly polarized negative surface is maintained only by mt-SAF19, which is also the paralog exhibiting the highest primary sequence and structural identity to human p32 (Fig. 5A). The peripheral edges of the respective side of the predicted Tb927.11.9600 trimer are also charged, while its central part is shielded by paralog-specific sequence extensions.

The remaining *T. brucei* p32 paralogs, Tb927.7.3470 and Tb927.11.9600, are not present in the assemblosome. However, when we performed IP-MS identification of interacting partners of the newly identified trypanosomal homologs of human TFB1M and NOA1 (Table S1), the protein Tb927.7.3470 was highly enriched along a substantial subset of assemblosome components (Fig. 5B). Thus, this p32-like protein is likely also associated with pre-mtSSU. Together, these results demonstrate that the p32 family has undergone expansion in *T. brucei* and that its members function predominantly during mtSSU biogenesis.

### p32 proteins have eukaryotic origin and their ancestral function is linked to gene expression in mitochondria

To reconstruct the evolutionary history of the six p32 family proteins identified in trypanosomes, we searched for p32 homologs in the predicted proteomes of Euglenozoa, a group encompassing kinetoplastids (including trypanosomatids; 14 species analyzed), their sister lineage of free-living marine diplonemids (10 species), and the early branching euglenids (5 species). We found that kinetoplastids and diplonemids typically encode six and four p32 family members, respectively, whereas euglenids express only a single homolog. A phylogenetic tree reconstructed from all obtained sequences (Fig. S4, Supplementary Data 1) revealed that the six paralogs in *T. brucei* were already present in the common ancestor of kinetoplastids.

The four diplonemid paralogs also cluster into well supported branches, however, the phylogenetic signal is insufficient to resolve the specific orthologous relationships between individual kinetoplastid and diplonemid paralogs (see legend of Fig. S4 for details). Three of the four p32-family proteins encoded by *Paradiplonema papillatum* were previously shown to copurify with core mtSSU components (12), indicating their role in mtSSU assembly in diplonemids. Neither the biochemical study nor our current search in genomic data identified an ortholog of mt-SAF24 in diplonemids. Together with the absence of U-indel editing, this suggests that a functional analog to p22 in *T. brucei* is not present in this group. Whether the diplonemid p32 proteins act as trypanosomal mt-SAF16/19/25 or differently cannot be inferred from the current analyses.

The presence of six p32 paralogs in trypanosomatids, in contrast to only single in yeast, humans and euglenids, prompted us to map the phylogenetic distribution of p32 family members across eukaryotes. We searched predicted proteomes of 196 species from all major eukaryotic groups and identified p32 homologs in all three main branches of eukaryotes, Discoba, Diaphoretickes, and Opimoda (Fig. 5C, Table S2). Approximately half of the analyzed organisms, including most opisthokonts, comprising fungi and humans, encode a single p32 protein. However, independent duplications have occurred frequently across divergent lineages. By far the largest expansion was detected within the eudicot plants, which encode 16 paralogs in *Arabidopsis thaliana* and 12 in *Populus trichocarpa*, strongly implying a significant functional diversification of p32 roles in plants.

Conversely, we did not detect any p32 in several major lineages, including the red algae (Rhodophyta), haptophytes, heteroloboseans, and jakobids (Fig. 5C, Table S2). Thus, the last eukaryotic common ancestor most likely contained a single ancestral p32 gene, which subsequently underwent independent duplications or was lost across different lineages.

Crucially, we found no evidence for the existence of the p32 family in Bacteria and Archaea, indicating that the family is an evolutionary invention of eukaryotes. Virtually all identified p32 proteins contain predicted N-terminal mitochondrial targeting signals (Table S2). Notably, p32 is absent from all organisms without mitochondrial genomes regardless their phylogenetic position (*Gregarina*, Metamonada, Archamoebea, *Blastocystis*, Breviata), strongly suggesting that the ancestral function of the protein is associated with mitochondrial genome expression.

## DISCUSSION

Biogenesis of mitochondrial ribosomes requires many transiently acting assembly factors (15,16). Structural studies have so far identified 38 factors acting in the assembly pathway of the exceptionally divergent mtSSU in *Trypanosoma brucei*, some of which are shared with bacteria or other eukaryotes (8,11,21). In this study, we identified a previously unrecognized trypanosomal mtSSU assembly factor, p22. This protein forms a stable homotrimer that caps a protrusion of the mtSSU precursor complex, the assemblosome.

Previously, p22 was reported as a protein required for the efficient U-indel RNA editing of the *COII* transcript, but not other mitochondrial mRNAs (24,58). We recapitulated these findings and further demonstrated that the steady-state levels of the core subunits of the RNA editing machinery remain unchanged upon p22 ablation (Fig. 2), confirming that the defect is specific to *COII* rather than a reflection of general editing impairment. While all other edited mitochondrial transcripts require *trans*-acting guide RNA molecules to define editing sites (65), *COII* editing, which involves insertion of four uridines (2), is guided by a sequence at the 3’ untranslated region (UTR) of the mRNA itself (66). This *cis*-editing mechanism requires a dedicated RECC module containing the endonuclease KREN3 (67). The editing defects in the p22 RNAi cell lines resemble those observed after KREN3 ablation (67). Possibly, p22 is involved in the inter-molecular interaction between the 3’ UTR and the editing site, or recruitment of the specialized RECC. Furthermore, the observed decrease in pre-edited *COII* mRNA levels suggests that p22 may also contribute to stabilization of the pre-mRNA.

The protein encoded by *COII* is a core membrane component of cIV. Therefore, as expected, abundance of cIV is decreased upon the induction of p22 RNAi. However, knockdown of p22 also impairs levels of ATP synthase and cIII. Each of these OXPHOS complexes contain a single mitochondrial encoded subunit, but editing of their transcripts is not severely affected upon ablation of p22 (24). Thus, defects of RNA editing associated with the loss of p22 cannot directly explain the decreased abundance of these complexes. The decreased levels of these complexes can be explained by secondary effects, for example by the mutual stabilization of individual complexes co-occurring in canonical cIII_2_cIV_2_ and unique cIIcIV_2_cV supercomplexes, both recently reported in *T. brucei* (68).

More plausibly, however, our data suggest that the drop in OXPHOS complexes in the absence of p22 stems from generally reduced efficiency of mitochondrial translation, caused by a defect in mtSSU assembly (Figs. 3&4). Supporting this scenario, nearly identical phenotypes are observed upon the downregulation of mt-SAF24, an essential mtSSU assembly factor with no functional or physical links to RNA editing machinery. Nevertheless, we note that primary defects in mitochondrial RNA editing and translation are inherently difficult to disentangle experimentally. The final phenotypic readout, observed as a collapse of OXPHOS complexes, loss of mitochondrial membrane potential, and compromised cell viability, likely reflects the dual role of p22 in both processes. Whether the functions of p22 in *cis*-editing and mitoribosome biogenesis are entirely independent or functionally or structurally coupled remains to be elucidated. Speculatively, p22 might mediate interplay between *COII* transcript maturation and mtSSU assembly to orchestrate OXPHOS biogenesis.

Protein p22 belongs to the p32 family. Mammalian p32, the prototypical representative of this family, was originally identified as an interacting partner of the nuclear mRNA splicing factor SF2/ASF (69). Since its discovery, it has been implicated in nuclear transcription, splicing, apoptosis, signaling and the biogenesis of both cytosolic and mitochondrial ribosomes. Mutations in human p32 are closely tied to severe metabolic pathologies, inflammation, and oncogenesis (reviews (70) and references therein). The protein has been detected in virtually all major cellular compartments, including the nucleus, cytosol, plasma membrane, extracellular secretory fractions, and mitochondria. While this multicompartmental localization and its “moonlighting” pleiotropy remain controversial in the literature, a consensus points to primary mitochondrial targeting (70–73). In murine mitochondria, p32 is indispensable for translation and co-immunoprecipitates multiple mitoribosomal proteins. It interacts with RNA, but not DNA, in vitro (74). Human p32 has been linked to mitochondrial rRNA processing (75). Based on three independent studies, human p32 interacts with the endonuclease YBEY to facilitate recruitment of mitoribosomal protein uS11m during mtSSU biogenesis (76–78). Mam33, the p32 homolog in *S. cerevisiae*, is a mitochondrial protein (79) required for the efficient translation of the cytochrome c oxidase subunit 1 transcript on fermentable carbon sources (80). Mam33 was shown to participate in mitoribosome biogenesis, but the biochemical evidence indicated its role in the mtLSU rather than mtSSU assembly (81).

Our findings implicating trypanosomal p22 in mtSSU biogenesis are therefore consistent with observations in the model organisms. However, unlike mammals and fungi, which express only single p32 protein, trypanosomatids encode an expanded family of six distinct paralogs. Five of its members are now directly implicated in the mtSSU assembly. The sixth paralog (Tb927.11.9600) is also confirmed to be mitochondrial by a high-throughput localization study (82), associating with the inner mitochondrial membrane as documented by subcellular proteomics (83) and submitochondrial fractionation (63). Remarkably, a paralog of *T. brucei* p22 and human p32 termed mS105 was identified as a component of the mature mtSSU in the green alga *Chlamydomonas reinhardtii* (84). Mirroring p22, mS105 assembles into a homotrimer (Fig. 5A); however, it associates with a lineage-specific rRNA extension located at a position entirely distinct from the sites occupied by the p22 homotrimer and the mt-SAF16/19/25 heterotrimer within the *T. brucei* assemblosome (Fig. 5D). Therefore, its structural role within the mature mitoribosome likely evolved independently from the transient assembly functions characterizing the trypanosomatid paralogs. Notably, neither the four *T. brucei* p32 paralogs present in the assemblosome nor the algal mS105 localize to the vicinity of uS11m (Fig. 5D). This strongly suggests that, in contrast to human p32, these p32 homologs are not involved in uS11m binding and they play a distinct role in mtSSU biogenesis or translation.

We mapped phylogenetic distribution of p32 proteins and did not detect any similar proteins in bacteria and archaea, implying that the family is specific for eukaryotes. Within eukaryotes, several groups lack p32 completely, but from the presence of the protein in all main branches it can be confidently inferred that p32 was present in LECA. After radiation of eukaryotes, the p32 family expanded in several lineages. The highest number of p32 paralogs is found in plants (16 paralogs in *A. thaliana*), followed by trypanosomatids, studied here (6 paralogs). This expansion of the p32 family in these two groups closely mirrors the evolutionary behavior of pentatricopeptide repeat (PPR) proteins. While present across eukaryotes, PPR proteins are massively multiplied in plants, where they participate in highly complex post-transcriptional processing in organelles, including RNA splicing, C-to-U editing, transcript stabilization, and mitoribosome assembly (85–87). Analogously to plants, trypanosomatids also feature an increased number of PPR proteins, which act in mitochondrial gene expression, including RNA editing, and contribute to mitoribosome assembly and architecture (88,89). Apparently, intricate gene expression pathways that evolved independently in these organisms require an expanded toolkit of proteins from p32 and PPR families.

On the other hand, p32 is absent from several eukaryotic groups. Most remarkably, we observed that every single organism devoid of a mitochondrial genome has lost p32. This correlation strongly supports the conclusion that the ancestral function of the p32 family, which emerged during eukaryogenesis, is tied to the expression of the mitochondrial genome. The clear structural or functional association of p32 family members with the mtSSU or its precursors across three deeply diverging eukaryotic lineages (90), represented by mammals, trypanosomatids, and green algae, indicates that a role in mitochondrial translation is a widespread property of the family. Nonetheless, the precise structural positioning, transient versus structural incorporation, and macromolecular interaction networks of these individual representatives differ substantially. Therefore, the specific ancestral function of p32 in mitochondrial translation apparatus remains to be elucidated.

## DATA AVAILABILITY

All data from this study are available in the manuscript or as Supplementary Data. The mass spectrometry proteomics data have been deposited to the ProteomeXchange Consortium via the PRIDE (91) partner repository with the dataset identifier PXD081257.

## SUPPLEMENTARY DATA

Supplementary Data are available at *NAR* Online.

## ACKNOWLEDGEMENTS

We thank Marek Vrbacký for processing, analyses and deposition of mass spectrometry proteomics samples and datasets, Laurie Read for kindly providing the anti-p22 antibody, and Jack D. Sunter for pJ1339.

## AUTHOR CONTRIBUTIONS

Conceptualization: P.C., O.G. Methodology: P.C., I.Š.S., J.E.W., J.Ř., A.Z., O.G. Formal Analysis: P.C., J.Ř. Investigation: P.C., I.Š.S., J.E.W, J.Ř., J.D., A.H., L.M., E.G. Resources: A.Z. Writing – Original Draft: P.C., O.G. Writing – review & editing: P.C., I.Š.S., A.Z., V.Y., O.G. Visualization: P.C., J.Ř., O.G. Funding acquisition: P.C., I.Š.S., J.E.W., V.Y., O.G.

