## Supplementary Information for "A p32 family RNA editing factor acts in mitochondrial ribosome biogenesis"

**List of Supplementary Data:**

**Supplementary Figures S1 to S4**

**Supplementary Table S1**: List of oligonucleotides, antibodies, *T. brucei* cell lines, reagents and chemicals, software, instruments and sequences.

**Supplementary Table S2**: IP-MS data

**Supplementary Table S3**: Distribution of p32 paralogs in species and higher taxa across eukaryotes including sequences and DeepLoc targeting prediction

**Supplementary Data 1**: A zip file with sequences of p32 in Euglenozoa and tree files (related to Supplementary Figure S4)

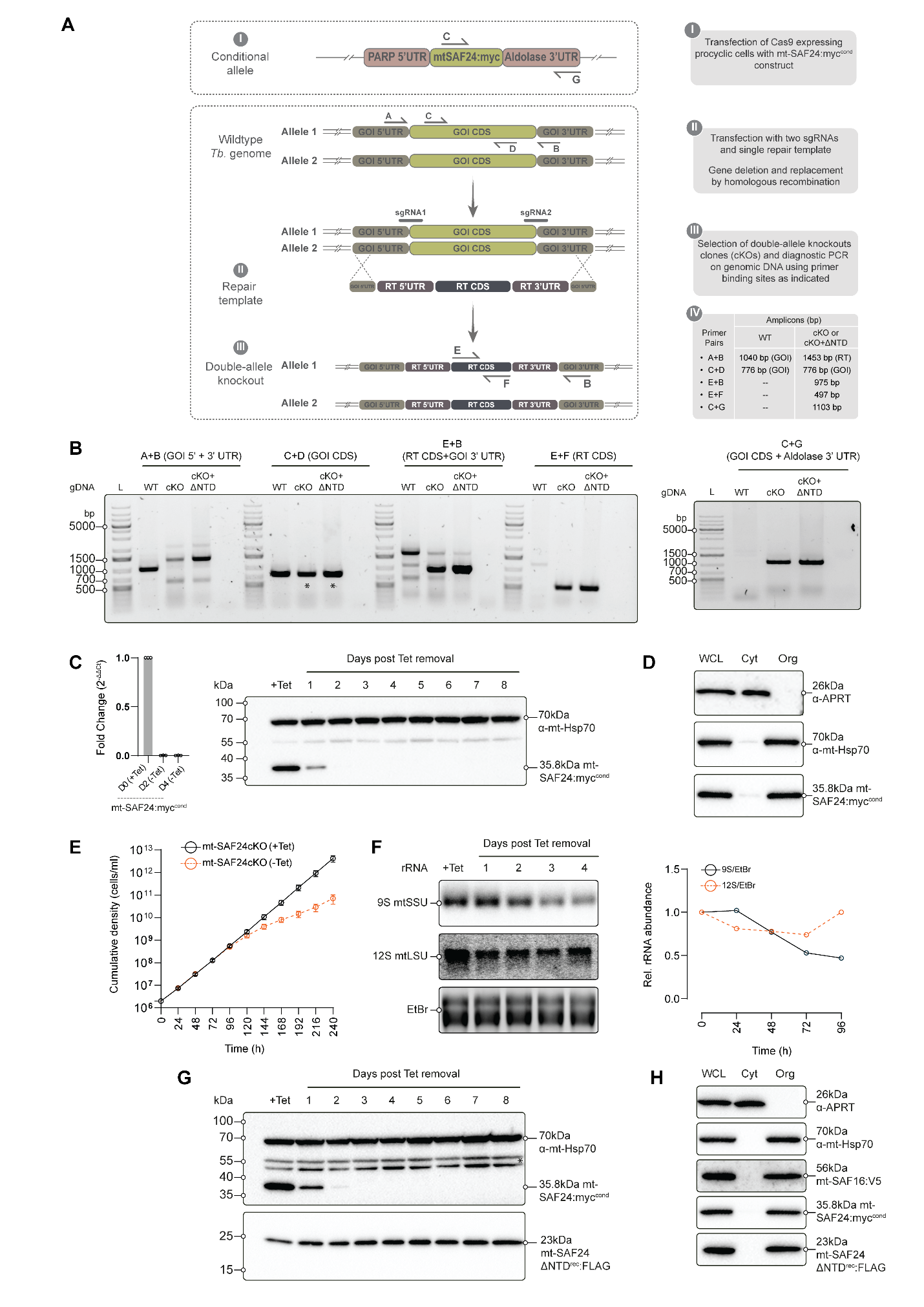

**Supplementary Figure S1. Genetic modifications of mtSSU assembly factors and their verifications. (A)** A scheme showing genetic modifications to knockout both mt-SAF24 alleles and positions of primers to verify the knock-out and the introduction of a conditional full-length myc-tagged mt-SAF24 (mt-SAF24:myc^cond^) in the resulting mt-SAF24^cKO^ strain. Lengths of amplicons expected with indicated primer pairs in wild-type (WT) and knock-out (KO) strains are shown. RT; repair template. **(B)** PCR verification of the mt-SAF24^cKO^ strain (KO) and a derived strain co-expressing mt-SAF24-ΔNTD^rec^:FLAG (KO+ΔNTD). **(C)** Tetracycline-regulated expression of mt-SAF24:myc^cond^ documented on RNA level by q-RT-PCR (values normalized to cells grown in the presence of tetracycline, β-tubulin gene used as a reference) and on protein level by immunoblotting with anti-myc and anti-mt-Hsp70 (loading control) antibodies. **(D)** Immunoblot of subcellular fractions showing organellar localization of mt-SAF24:myc^cond^ (Wcl - whole cell lysate; Cyt - cytosolic fraction; Org - organellar fraction containing mitochondria). **(E)** Growth curve of the mt-SAF24^cKO^ cell line in presence of tetracycline and after tetracycline removal. **(F)** Northern blot analysis of mtSSU (9S) and mtLSU (12S) rRNA levels in the mt-SAF24^cKO^ cells at indicated time points after tetracycline removal. The signal from ethidium bromide-stained gel prior transfer was used as a reference for quantification shown in the right panel. **(G)** Western blot documenting tetracycline-regulated expression of mt-SAF24:myc^cond^ and constitutive expression of mt-SAF24-ΔNTD^rec^:FLAG with anti-myc, anit-FLAG and anti-mt-Hsp70 (loading control) antibodies. **(H)** Immunoblot of subcellular fractions showing organellar localization of mt-SAF24:myc^cond^, mt-SAF24-ΔNTD^rec^:FLAG and mt-SAF16:3×V5.

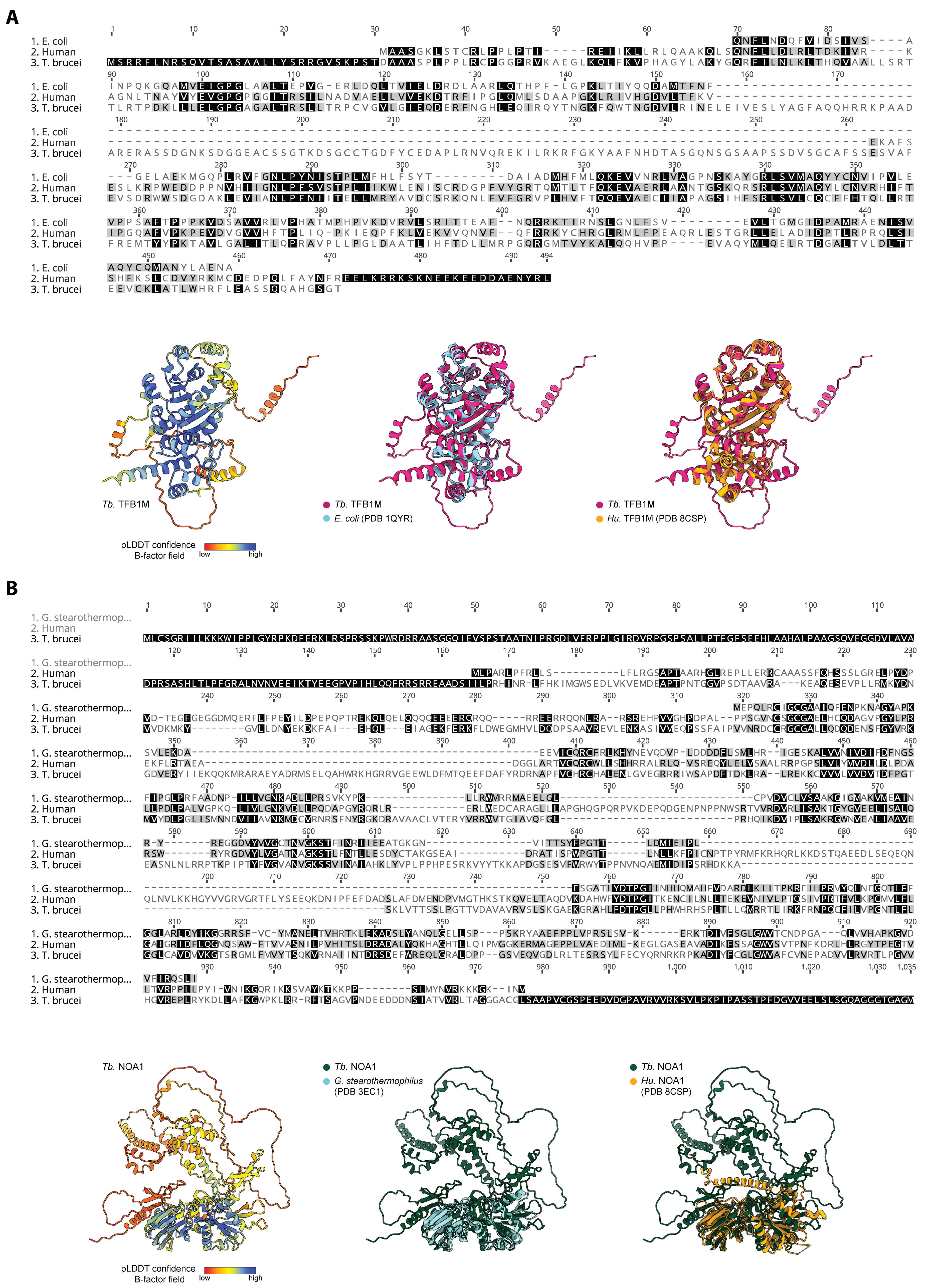

**Supplementary Figure S2. *Trypanosoma brucei* encodes conserved assembly factors TFB1M and NOA1. (A)** Sequence and structural alignments of *Escherichia coli* KsgA (1), human (2) and *T. brucei* homologs of TFB1M. **(B)** Sequence and structural alignments of *Geobacillus stearothermophilus* YqeH (3), human (2) and *T. brucei* homologs of NOA1. Structures of *T. brucei* proteins were predicted by AlphaFold 3 (4). All other structures were determined experimentally, and PDB accession numbers are included. Sequence alignments were constructed and visualized in Geneious.

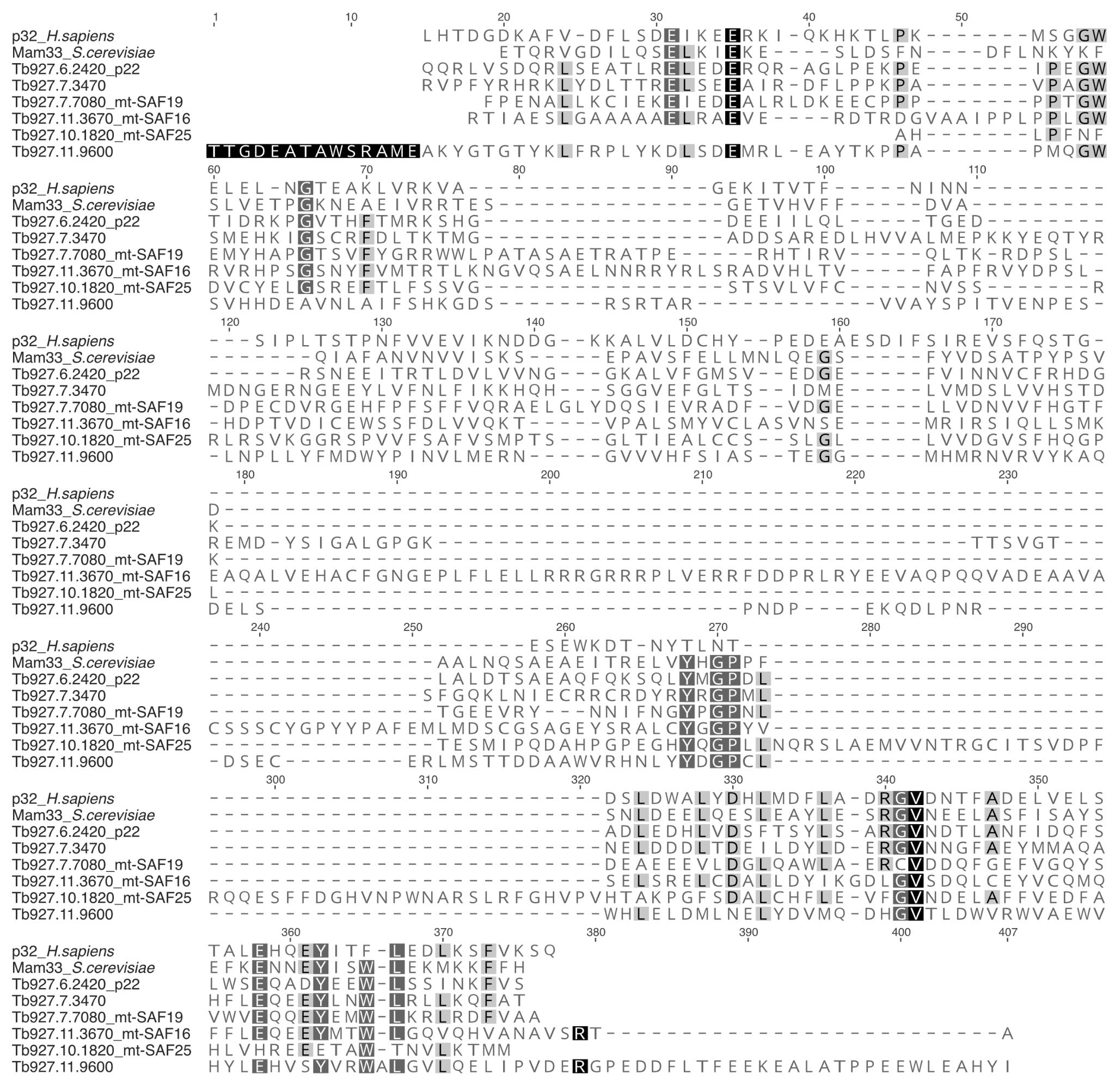

**Supplementary Figure S3. Sequence alignment of p32 family proteins.** Human p32, Mam33 from *Saccharomyces cerevisiae*, and all six p32 paralogs from *T. brucei* were aligned based on their structures using PROMALS3D, the alignments were visualized in Geneious, adjusted manually, N-terminal predicted mitochondrial sequences or unstructured regions were removed.

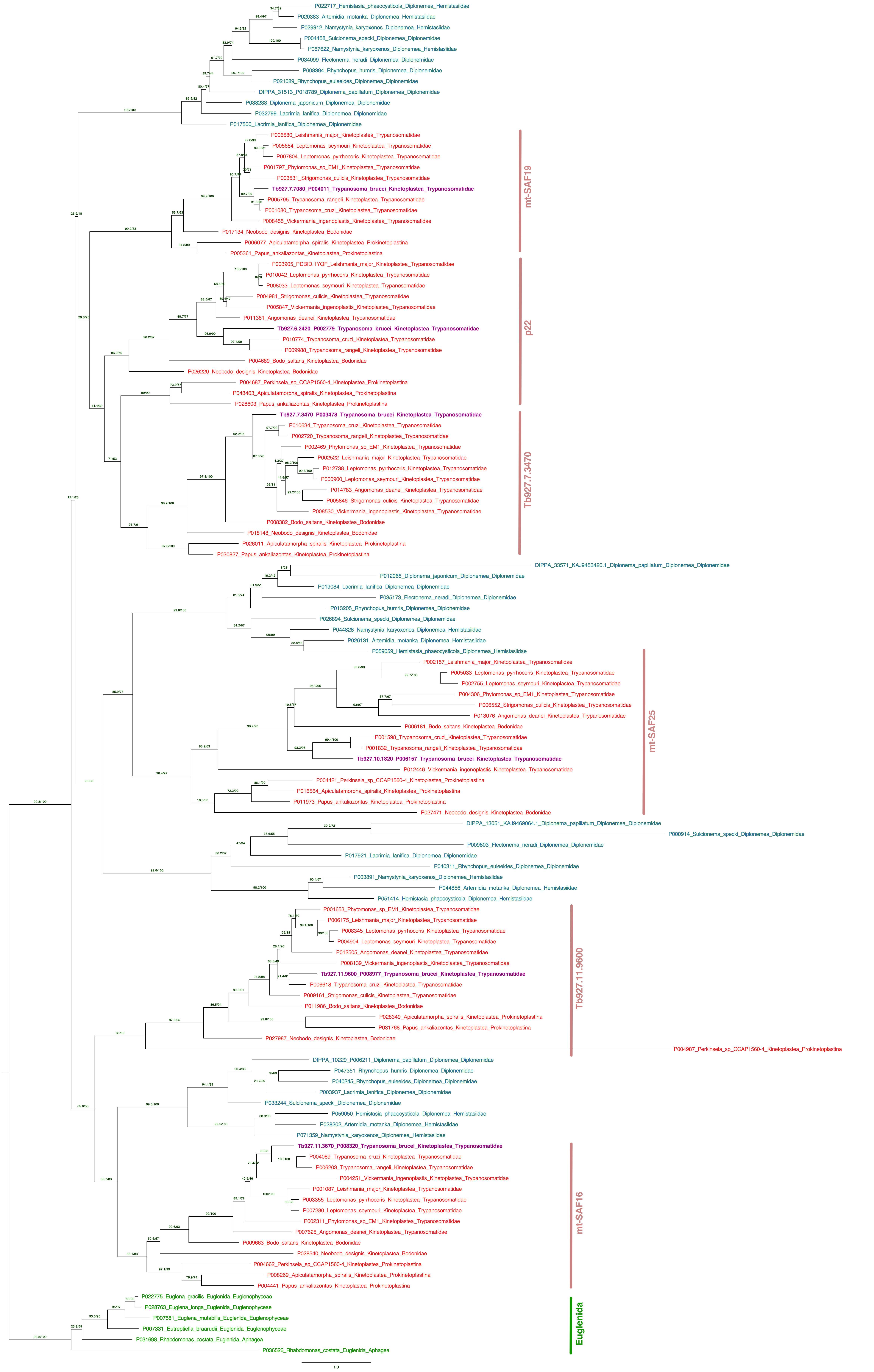

**Supplementary Figure S4: Phylogenetic tree of p32 proteins from Euglenozoa.** The branches corresponding to orthologs of individual *T. brucei* p32 proteins in kinetoplastids and the branch of the single p32 protein in euglenids are indicated. The remaining proteins (shown in blue) are from diplonemids. The tree was visualized in FigTree v1.4.4 ( <https://tree.bio.ed.ac.uk/software/figtree/>).
